# Detecting Somatic Mutations Without Matched Normal Samples Using Long Reads

**DOI:** 10.1101/2024.02.26.582089

**Authors:** Jared T. Simpson

## Abstract

DNA sequencing of tumours to identify somatic mutations has become a critical tool to guide the type of treatment given to cancer patients. The gold standard for mutation calling is comparing sequencing data from the tumour to a matched normal sample to avoid mis-classifying inherited SNPs as mutations. This procedure works extremely well, but in certain situations only a tumour sample is available. While approaches have been developed to find mutations without a matched normal, they have limited accuracy or require specific types of input data (e.g. ultra-deep sequencing). Here we explore the application of single molecule long read sequencing to calling somatic mutations without matched normal samples. We develop a simple theoretical framework to show how haplotype phasing is an important source of information for determining whether a variant is a somatic mutation. We then use simulations to assess the range of experimental parameters (tumour purity, sequencing depth) where this approach is effective. These ideas are developed into a prototype somatic mutation caller, smrest, and its use is demonstrated on two highly mutated cancer cell lines. Finally, we argue that this approach has potential to measure clinically important biomarkers that are based on the genome-wide distribution of mutations: tumour mutation burden and mutation signatures.

## 1 Introduction

Large scale efforts to sequence thousands of cancer genomes has been a major undertaking since the invention of high-throughput DNA sequencing instruments. These projects have catalogued driver mutations (Weinstein et al. 2013; Bailey et al. 2018; Rheinbay et al. 2020; “Pan-cancer analysis of whole genomes” 2020), defined mutational processes and the signatures they leave on the genome (Alexandrov, Nik-Zainal, et al. 2013; Alexandrov, Kim, et al. 2020; Nik-Zainal et al. 2012), tracked the evolutionary trajectories of tumours (Shah et al. 2009; Sottoriva, Spiteri, et al. 2013; Sottoriva, Kang, et al. 2015; Gerstung et al. 2020), uncovered the originating cell and tissue types of tumours (Jiao et al. 2020; Hendrikse et al. 2022) and discovered mutation-based biomarkers that can guide treatment choice (Chan et al. 2019; Zhao et al. 2019; André et al. 2020). Underpinning these studies was the development of both the high throughput sequencing instruments and analysis methods that can accurately detect mutated bases from the sequenced reads. Unlike in typical human genome sequencing projects, where one typically wants to discover variation within a sequenced genome compared to a reference genome, cancer genome projects aim to discover *somatic* mutations that occurred during the development and progression of a tumour. This requires finding genetic variation within an individual, where a subset of cells contain a mutated copy of a chromosome with respect to the chromosome inherited from the individual’s parents (see **Figure1a**).

**Figure 1:**
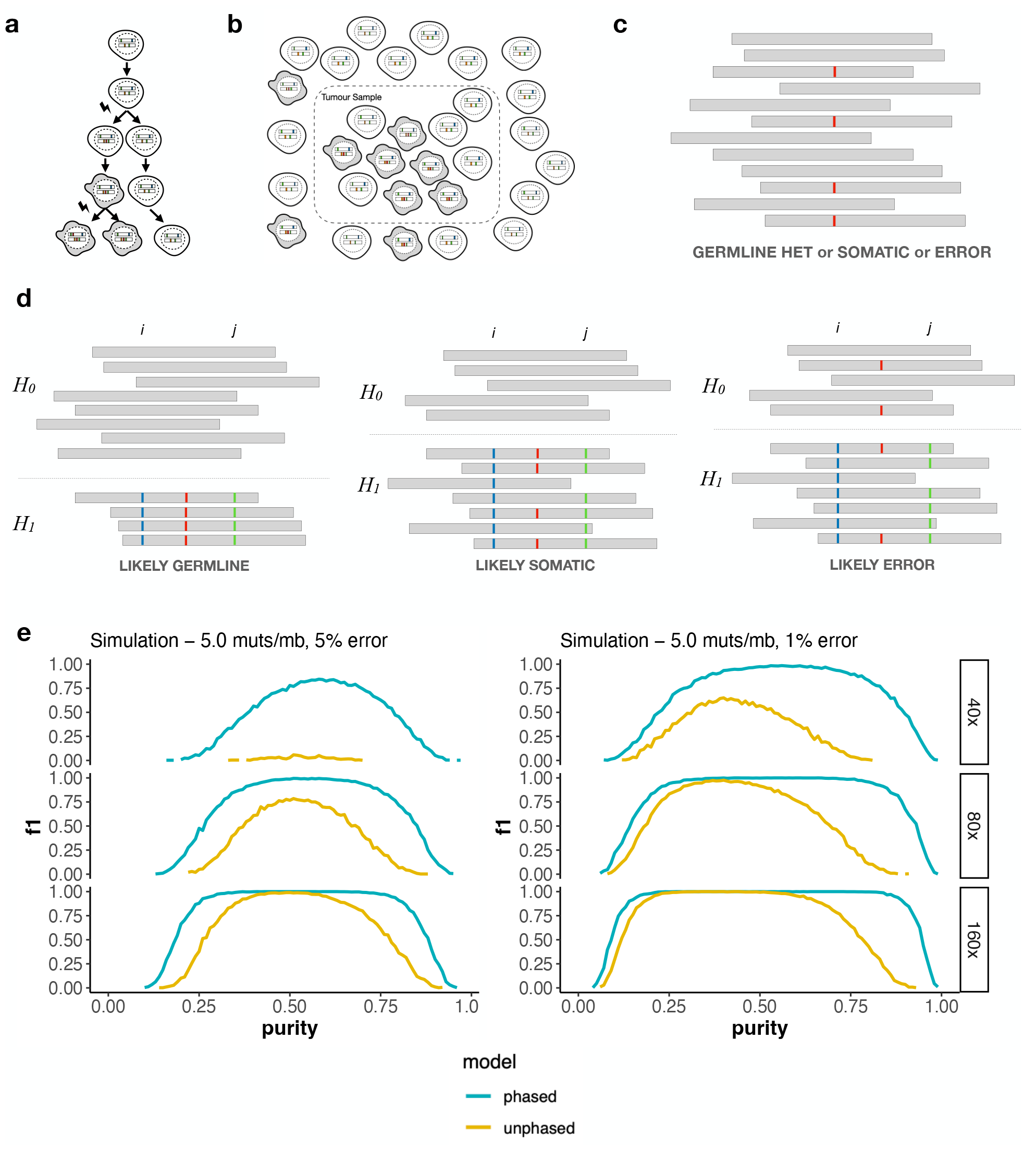
**a**. Cell lineages accumulate somatic mutations over time. When a healthy cell (white) becomes malignant (grey) the ns contained within that lineage rise in frequency. Additional mutations within the tumour population may generate es, where the mutations are contained in a subset of the tumour cell population. **b**. Tumour samples are typically a of cancerous cells and normal cells. The proportion of tumour cells is called the tumour *purity* and denoted *α*. When ng DNA from the tumour sample the somatic mutations will present as mosaics with only some of the sequenced reads ing the mutation. **c**. Variant callers that do not use phasing information need to make a decision based on the number of rey bars) that support the alternate allele (red) and reference allele. Depending on the number of reads supporting the allele it may be ambiguous whether the position is a heterozygous SNP, sequencing error or somatic mutation. **d**. If otype of each read (*H*_0_, *H*_1_) can be determined using nearby heterozygous SNPs (positions *i* and *j* with blue and green, respectively) the arrangement of reads supporting the alternate allele (red) can help determine whether it is a somatic n. We expect all phased reads to support the alternate allele when the position is a heterozygous SNP (left) or to support re of reference and alternate alleles when it is a somatic mutation (middle). When the position contains a sequencing do not expect the evidence of the error to segragate by haplotype (right). **e**. Simulation results demonstrating that plotype phasing information can classify somatic mutations more accurately across a range of tumour purity, read depth r rates.

The gold standard method of detecting somatic mutations is to compare sequencing data from the tumour to a matched “normal” sample that is assumed to be representative of the individual’s inherited genome. This method, referred to as tumour-normal calling, is highly accurate and widely used (Koboldt et al. 2009; Larson et al. 2012; Saunders et al. 2012; Cibulskis et al. 2013; Fang et al. 2021). Tumour-only calling methods have also been developed for situations where a normal sample is not available (e.g. for biobanked tissue samples where a normal was not collected, or to simplify clinical workflows). These methods typically rely on analysis of the fraction of reads supporting a candidate mutation, as this may differ from the fraction supporting inherited heterozygous SNPs, usually coupled with extensive filtering of the candidate calls against databases of variants known to occur in the human population (Smith et al. 2016; Kalatskaya et al. 2017; Sun et al. 2018). While this strategy can be effective in certain situations, like intermediate purity tumours that are very deeply sequenced (Sun et al. 2018), it is inherently less accurate than tumour-normal pair calling, and filtering against population database raises issues of bias for underrepresented populations (Nassar et al. 2022).

Thus far, cancer genome sequencing projects have primarily relied on highly accurate short read sequencing technologies. Long read sequencing technologies from Oxford Nanopore Technologies (ONT) and Pacific Biosciences (PacBio) are increasingly accurate (Sereika et al. 2022; Kolesnikov et al. 2023), and improvements to instrument throughput have expanded the range of possible applications. Long read sequencing is now the gold standard for genome assembly (Rhie et al. 2021; Nurk et al. 2022). Both ONT and PacBio sequencers can measure the genome and epigenome simultaneously by directly detecting base modifications (Flusberg et al. 2010; Laszlo et al. 2013; Schreiber et al. 2013; Simpson et al. 2017). Recently, a tumour-normal calling approach has been developed for long reads that has comparable accuracy to short read sequencing (Zheng, Su, et al. 2023).

In this paper, we explore the potential for using long read sequencing to perform tumour-only mutation calling. The advantage of long reads for this problem is that reads can often be assigned to individual haplotypes, transforming the problem of detecting somatic mutations from potentially small shifts in the variant allele fraction, into detecting the presence of two or more bases within a single haplotype, as proposed in the mosaic variant detection method by Darby et al. (2019) for 10X Genomics linked reads. In this work we formalize the problem and use simulations to assess the applicability of this approach as a function of key experimental parameters (sequencing depth, tumour purity, sequencing error rate). Then, we develop a prototype mutation caller, *smrest* (somatic mutation rate estimator), for real long read data and demonstrate its use on cancer cell lines. Finally we present the intended application of this tool, which is to discover genome-wide mutation patterns that can be used to guide therapy choice, such as tumour mutation burden or mutation signatures.

## 2 Results

### 2.1 Overview of Method and Feasibility

Somatic mutations are by definition mosaic; they occur in a subset of cells within the human body. When a particular cell becomes cancerous the complement of somatic mutations contained within that originating cell, and subsequent mutations that occur during the tumour’s expansion, rise in frequency (**Figure 1a**). These mutations can be detected by comparing DNA sequences from a tumour sample to a blood or non-cancerous tissue from the same individual (tumour-normal calling, reviewed in Xu 2018). The subject of this paper however is calling mutations in absence of the matched normal sample, referred to as “tumour-only” calling. This problem has been studied for short read sequencing and, most relevant to this work, Darby et al. 2019 introduced the idea of using haplotype phasing patterns for 10X Genomics Linked Reads.

Importantly for these prior approaches, and essential to this work, is that real tumour samples are typically mixtures of both cancerous and healthy cells (**Figure 1b**). The fraction of cancerous cells is referred to as tumour purity or tumour cellularity which we will denote *α*. The presence of normal cells provides evidence of the allele that the individual inherited from their parent. In a sequencing experiment this may shift the variant allele fraction (VAF; the proportion of reads supporting the putative mutation) away from the ratio expected of heterozygous SNPs (0.5 for copy number balanced autosomes, as each parental haplotype is equally likely to be sampled by a read). The ability to detect mutations from the shift in VAF alone is strongly dependent on sequencing depth, sequencing error rate and tumour purity (**Figure 1c**). For example, when tumours are nearly pure (*α ≈* 1) somatic mutations are nearly indistinguishable from heterozygous SNPs and at the other extreme, where few cells in the sample derive from the tumour (*α ≈* 0), somatic mutations will look like sequencing errors. If the parental haplotype (maternal or paternal) of each read is known, then the problem of identifying somatic mutations simplifies to identifying whether there is sufficient evidence of a mixture of alleles within a haplotype (one inherited allele from the normal cells, and the mutant allele from the cancerous cells). **Figure 1d** illustrates how phased reads bearing evidence of a variant can help identify whether it is a heterozygous SNP, somatic mutation or sequencing error.

To explore the feasibility of somatic mutation calling from long read sequencing we developed statistical models for phased and unphased data to calculate the posterior probability that a given position of the genome harbors a somatic mutation based on the number of reads supporting the reference and alternate allele, along with the tumour’s purity and sequencing error rate (see **Methods**). We then generated simulated data according to this model to explore the relationship between sequencing depth, error rate and purity (**Figure 1e**). In all parameter sets tested haplotype-phased data provides equal or superior variant calling accuracy. For a fixed depth and sequencing error rate this enables accurate somatic mutation calling across a wider range of tumour purity. For example, at 80x sequencing depth with 1% error rate the somatic mutation calling F1 exceeded 0.9 for simulated samples with purity in the range 0.23 *−* 0.84 when the data was phased but only 0.42 *−* 0.52 when the data is not phased.

### 2.2 Mutation Calling on Single Molecule Long Reads

The simulations presented above demonstrate that haplotype phasing can improve somatic mutation calling accuracy. These simulations assume ideal data however, where every read can be correctly assigned to a haplotype and the reads are perfectly aligned to the reference genome. As a further proof of principle, we developed and benchmarked a mutation caller, called *smrest* (somatic mutation rate estimator) for real data. Briefly, this program first detects heterozygous SNPs, phases them, partitions reads into haplotypes (haplotagging), then calls mutations using a procedure that extends the simulation model with a probabilistic alignment framework that is widely used in other variant callers (Albers et al. 2011; H. Li 2011; Garrison and Marth 2012; Poplin et al. 2018; Cooke, Wedge, and Lunter 2021). Finally, putative somatic variants are filtered to remove likely artifacts (e.g. from mapping errors, or systematic sequencing errors) and the resulting mutations that pass quality control are output as a VCF file. The mutation caller also generates a BED file describing the regions of the genome that were deemed callable (both haplotypes detected with sufficient sequencing depth). A high-level presentation of the mutation calling procedure is provided here, with complete details in the **Methods**.

#### Genotyping and Phasing

The input to standard haplotype phasing algorithms is a set of heterozygous SNPs (Patterson et al. 2015). While variant calling and genotyping long read data from human genomes can now be done with high accuracy (Zheng, S. Li, et al. 2022) cancer genomes may have copy number imbalances that shift the read support of heterozygous SNPs away from the expectation of equal support for the paternal and maternal alleles. In the worst case where one parental haplotype is entirely lost, known as loss-of-heterozygosity (LOH), the evidence of the lost allele will only come from the normal cells within a sample and hence be a function of the tumour’s purity. To account for these factors we developed a genotyper for known variant sites that relaxes the assumption of copy number balance. This method is designed to be conservative and prefer false negatives over false positives, as the latter is more likely to introduce erroneous haplotype assignments. The set of heterozygous SNPs found by this procedure is used as input to `whatshap` (Patterson et al. 2015) for phasing.

#### Mutation Calling and Filtering

The phased VCF file, and the original BAM file containing the reads mapped to the human reference genome, are input into the mutation calling program `smrest_call`. This program haplotags each read, discovers candidate variants, then calculates class probabilities for each candidate using a read-haplotype likelihood model (see **Methods**). Finally, summary statistics are gathered for each output call to facilitate mutation filtering, e.g. when variants show evidence of strand bias.

### 2.3 Mutation Calling in COLO829

To characterize the performance of smrest for long read tumour-only somatic mutation calling we first analyzed COLO829, a melanoma cell line with a matched normal (COLO829BL). In all experiments we mixed reads from the tumour and normal together, without knowing the origin of each read, to simulate tumour-only sequencing with a controlled purity and sequencing depth. COLO829 has a high mutation rate of *≈*14 mutations per megabase (Titmuss et al. 2022) due to ultraviolet light damage (Pleasance et al. 2010), and is frequently used to benchmark the performance of sequencing technologies and analysis methods (Arora et al. 2019; Espejo Valle-Inclan et al. 2022). We downloaded Oxford Nanopore R10.4.1 reads for both COLO829 and COLO829BL from the Oxford Nanopore Open Datasets Collection. The tumour and normal sample were sequenced to 95x and 57x sequencing depth, respectively. The reads were prepared with a mixture of standard and ultra-long library preparations with 7.5x and 7.1x coverage of reads exceeding 100kbp in length, simplifying haplotype phasing. We also downloaded Illumina short reads for this pair of samples. Access information for all datasets used in this work are provided in **Table 1**.

**Table 1:** Summary of datasets used in this manuscript.

| Sample | Technology | Location/Accessions | Reference |
| --- | --- | --- | --- |
| COLO829 | ONT | colo829_2023.04/COLO829/ | - |
| COLO829BL | ONT | colo829_2023.04/COLO829BL/ | - |
| COLO829 | PacBio-HiFi | revio/2023Q2/COLO829 | - |
| COLO829BL | PacBio-HiFi | revio/2023Q2/COLO829-BL | - |
| COLO829 | Illumina | ERR2752450 | Cameron, Baber, Shale, Valle-Inclan, et al. 2021 |
| COLO829BL | Illumina | ERR2752449 | Cameron, Baber, Shale, Valle-Inclan, et al. 2021 |
| HCC1395 | ONT | SRR25005626 | Zheng, Su, et al. 2023 |
| HCC1395BL | ONT | SRR25005625 | Zheng, Su, et al. 2023 |
| HCC1395 | PacBio-HiFi | revio/2023Q2/HCC1395 | - |
| HCC1395BL | PacBio-HiFi | revio/2023Q2/HCC1395-BL | - |
| HCC1395 | Illumina | WGS_IL_T_1.bwa.dedup.bam | Xiao et al. 2021 |
| HCC1395BL | Illumina | WGS_IL_N_1.bwa.dedup.bam | Xiao et al. 2021 |
| CliveOME | ONT | cliveome_kit14_2022.05/ | - |

**Table 2:** Summary of software used in this manuscript.

| Software | Version | Reference |
| --- | --- | --- |
| minimap2 | 2.24-r1122 | H. Li 2018 |
| bwa | 0.7.17-r1188 | H. Li 2013 |
| samtools | 1.16 | H. Li et al. 2009 |
| bcftools | 1.16 | Danecek et al. 2021 |
| clairS | 0.1.16 | Zheng, Su, et al. 2023 |
| mutect2 | 4.4.0.0 | Benjamin et al. 2019 |
| whatshap | 1.7 | Patterson et al. 2015 |
| bedtools | 2.30.0 | Quinlan and Hall 2010 |
| snakemake | 7.19.1 | Köster and Rahmann 2018 |

To provide context to the performance of smrest we also ran ClairS (Zheng, Su, et al. 2023), a recently developed tumour-normal pair (abbreviated TN hereafter) caller based on neural networks on the long read data. For short reads we ran Mutect2 (Benjamin et al. 2019), which is designed for TN calling but also supports tumour-only (abbreviated TO) calling via filtering against population databases and a panel-of-normals to remove sequencing artifacts. It is included here to compare the performance of smrest against a short read method that is commonly used when a matched normal sample is not available.

To prepare the datasets both the tumour and normal cell line samples were downsampled to coverage of 20x or 40x. For the TO methods the read sets were merged together to give a single sample with equal coverage of the tumour and normal (50% purity). The tumour purity parameter to smrest was set to this known value (*α* = 0.5). For the TN callers the tumour and normal read sets were provided as separate inputs.

The mutation calls for all programs were compared to an externally curated call set (**Methods**) that was considered the ground truth. We discarded all mutations (in the truth data, or any call set from any program) with predicted VAF *<* 10% as low VAF mutations cannot be reliably called at the sequencing depths used for this analysis. In addition, we only considered substitution variants. As cell lines accumulate mutations (Petljak et al. 2019) that will appear as somatic mutations in tumour-only analysis we identified putative mutations in COLO829BL using the short read datasets and Mutect2 in TN mode by swapping the tumour and normal inputs, and filtering the results to avoid calling inherited SNPs as mutations in loss-of-heterozygosity regions (**Methods**). Any called mutations found in this COLO829BL mutation list were ignored for subsequent analysis. We characterized the accuracy of each program on each dataset in the regions of the genome designated as “high confidence” (denoted HC, **Figure 2a**), and the subset of the HC regions that could be successfully called and phased by smrest (**Figure 2b**) as precision-recall curves stratified by a variant’s QUAL score (for smrest and ClairS) or TLOD (for mutect2-TN and mutect2-TO).

**Figure 2:**
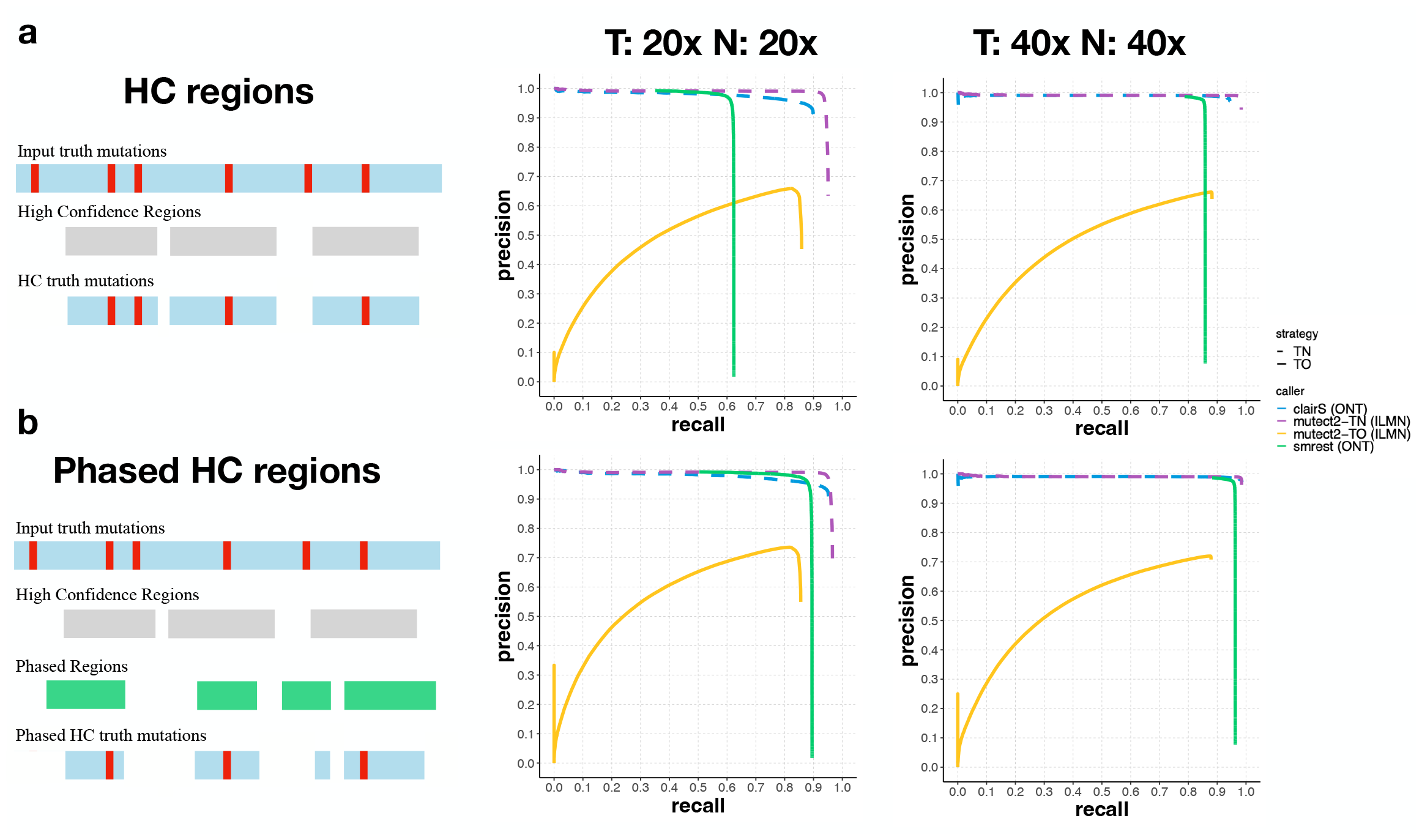
Somatic mutation calling performance on COLO829. Short and long read datasets were prepared with 20x tumour and 20x normal coverage and 40x tumour and 40x normal coverage. Each data set was provided as input to smrest (green solid line), clairS (blue dashed line), mutect2 in TN mode (purple dashed line) or mutect2 in TO mode (yellow solid line). Precision/recall curves were calculated using an external call set as ground truth. Two subsets of the mutation calls were analyzed, one consisting of calls that lie in regions deemed High Confidence (**Panel a**, top row) and one consisting of calls that lie in the HC regions that were successfully phased by smrest at the given coverage (**Panel b**, bottom row).

As expected the variant callers that use a tumour-normal strategy perform extremely well, even for the lowest depth dataset (20x/20x). For tumour-only calling the mutect2-based approach can achieve high recall, but with limited precision due to the difficulty of identifying somatic mutations from short reads even when provided with a database of population variants. In contrast, smrest achieves high precision without a matched normal and without filtering against population databases. This method is not a replacement for tumour-normal calling however. When the entire set of high-confidence regions is considered (spanning 2, 080Mbp of human reference GRCh38) recall ranges from 0.61 (20x/20x) to 0.86 (40x/40x). Sensitivity is primarily determined by the ability to phase the genome, as recall improves to 0.87-0.96 when only successfully phased high-confidence regions are considered. This highlights the critical need for genome-wide phasing if this approach is to be used for comprehensive somatic mutation detection.

Our method was primarily developed, tested, and debugged on COLO829 data sequenced with Oxford Nanopore long reads. To assess whether our approach generalizes, we additionally analyzed COLO829 PacBio HiFi data, and a highly mutated breast cancer cell line (HCC1395/HCC1395BL) sequenced with both ONT R10.4.1 and PacBio HiFi reads. The accuracy of each mutation call set was calculated as above, however here we computed the intersection of the ONT and PacBioHiFi phased regions to ensure all call sets were analyzed over the same regions of the genome. For COLO829, the Oxford Nanopore call set from smrest had slightly higher recall than the PacBio call set (**Figure 3a**), likely due to the presence of ultra-long reads that simplify the construction of long range haplotypes. When computing accuracy over the phased regions of the genome the ONT and PacBio data were both highly accurate. ClairS performed very well with data from each technology. The HCC1395/HCC1395BL cell line is more challenging than COLO829 due to the presence of a long tail of low VAF mutations (**Figure 3c,d**) possibly indicating multiple subclones. While smrest was able to precisely call mutations for both the ONT and PacBio datasets, the PacBio dataset had higher recall, likely due to the higher accuracy allowing lower frequency mutations to be identified. This trend is also seen in ClairS where the recall on PacBio data was somewhat higher than on ONT reads.

**Figure 3:**
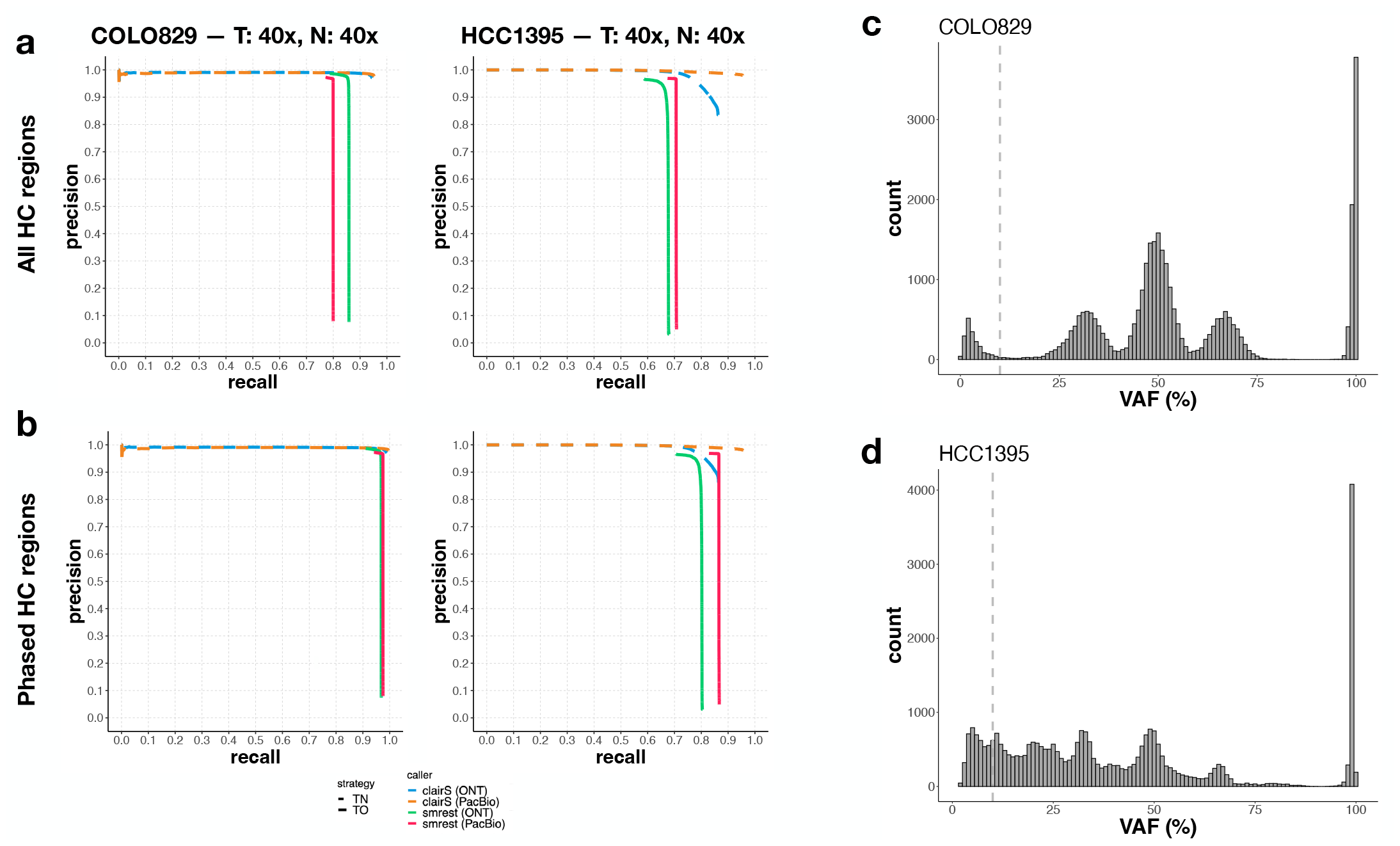
Somatic mutation calling performance on 40x tumour and normal coverage of COLO829 (left column of panel **a**. and **b**. and HCC1395 (right column) with ONT and PacBio data. As in the previous figure, callset accuracy was calculated over the High Confidence regions (**a**.), or the phased subset of the HC regions (**b**.). Each data set was provided as input to smrest (ONT: green solid line, PacBio: red solid line) or clairS (ONT: blue dashed line, PacBio: orange dashed line.). The distribution of variant allele frequencies for the truth call set for each sample is shown in (COLO829: **c**., HCC1395: **d**.), with the lower VAF threshold for inclusion within the analysis annotated as a vertical dashed line.

### 2.4 Assessing the Effect of Tumour Purity

In our simulations (**Figure 1e**) we observed that variant calling performance is a function of tumour purity. We therefore performed a series of experiments to assess this effect on the real datasets. Here we selected tumour purity in the range 0-100%, then computed the number of COLO829 (tumour) and COLO829BL (normal) reads needed to reach a total of 40x or 80x coverage at the selected purity. Each mutation calling program was run and analyzed as described above. For this analysis smrest was not provided the known value of tumour purity, it was left at its default value of *α* = 0.5.

In addition, only the phased HC regions were assessed. The results are shown in **Figure 4**. As expected and seen in the simulations, the performance of all mutation calling approaches is limited at extreme values of tumour purity as at low purity there is insufficient tumour depth to confidently identify mutations and at high purity there is insufficient normal depth to confidently say what the inherited allele is. Our program achieves comparable performance to the TN callers and in particular maintains high precision across a wide range of tumour purities, at both 40x and 80x coverage. In contrast, short read TO calling fails to achieve high precision. These results highlight the importance of sequencing depth as 80x coverage allows a much wider range of (simulated) sample purities to be reliably analyzed, as predicted by our simulations.

**Figure 4:**
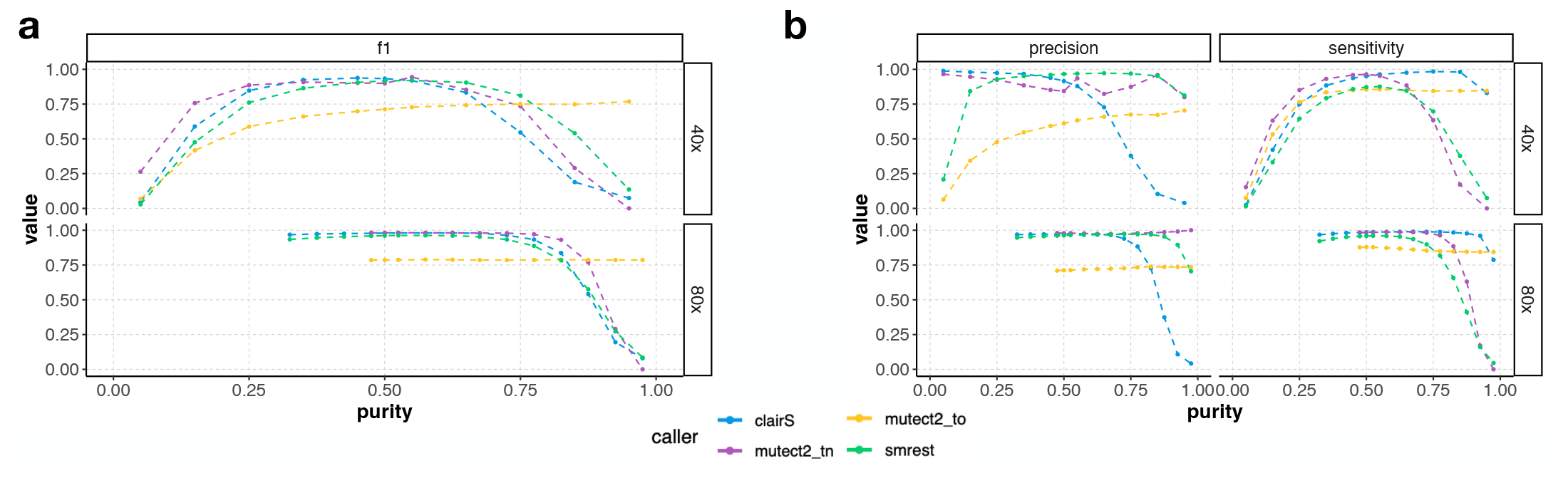
Somatic mutation calling accuracy (**Panel a**: F1, **Panel b**: precision and sensitivity) metrics as a function of tumour purity for 40x (top facet within each panel) and 80x (bottom) coverage sequencing of COLO829/COLO829BL with smrest (green), clairS (blue), mutect2-TN (purple) or mutect2-TO (yellow)

### 2.5 Estimating Tumour Mutation Burden and the Mutation Spectrum

Cancer genome sequencing is increasingly used to help guide treatment choices. While there are now many known point mutations, indels and structural variants that may indicate response or resistance to certain therapies (Krysiak et al. 2023), an emerging class of biomarker is based on the overall pattern of mutation across the genome. Perhaps most prominently, the tumour’s mutation burden (TMB; the number of observed coding mutations per megabase of coding sequence) is used for predicting response to immunotherapies, for example pembrolizumab (Marabelle et al. 2020) for tumours classified as TMB-high (*≥* 10 mutations/MB, Marcus et al. 2021). PARP inhibitors are used to treat patients with mutations in the homologous repair pathways (Patel, Sarkaria, and Kaufmann 2011; Miller et al. 2020), which can be detected with high accuracy using mutation signatures (Davies et al. 2017). As both TMB and mutation signatures are genome-wide phenomena they can be determined by sequencing only a subset of the genome (Milbury et al. 2022).

While we have demonstrated that in certain situations our approach can be highly sensitive, the requirement of phasing every base of the genome that might harbour a mutation of interest may limit the application of this approach for general purpose somatic mutation detection. Therefore our primary intended use case is to detect biomarkers that can be found from accurate mutation calling in defined regions of the genome. In our case, we propose to calculate TMB from the phased subset of the high confidence regions, by dividing the number of called mutations by the total size of the genome phased. To assess the feasibility of this strategy we performed additional mixture experiments where reads from COLO829 and COLO829BL were again merged together into a single sample with a defined purity (30%-70%, in steps in 10%) and depth (10x to 82x, in steps of 8x). Here we considered all mutations that passed our filtering criteria with a minimum quality score of 20 to be called. In this analysis we included a healthy blood sample openly released by Oxford Nanopore to assess the false positive rate in a sample that is assumed to be free from somatic mutations. In **Figure 5a**, we observe that the estimated mutation burden converges to the expected value of 14 mutations per megabase of analyzed sequenced (Titmuss et al. 2022), with the exception of the 30% purity sample which underestimates TMB at 13.1 muts/MB at the maximum analyzed depth of 74x. In the healthy blood sample few mutations were called (maximum of 0.2 muts/MB at 66x).

**Figure 5:**
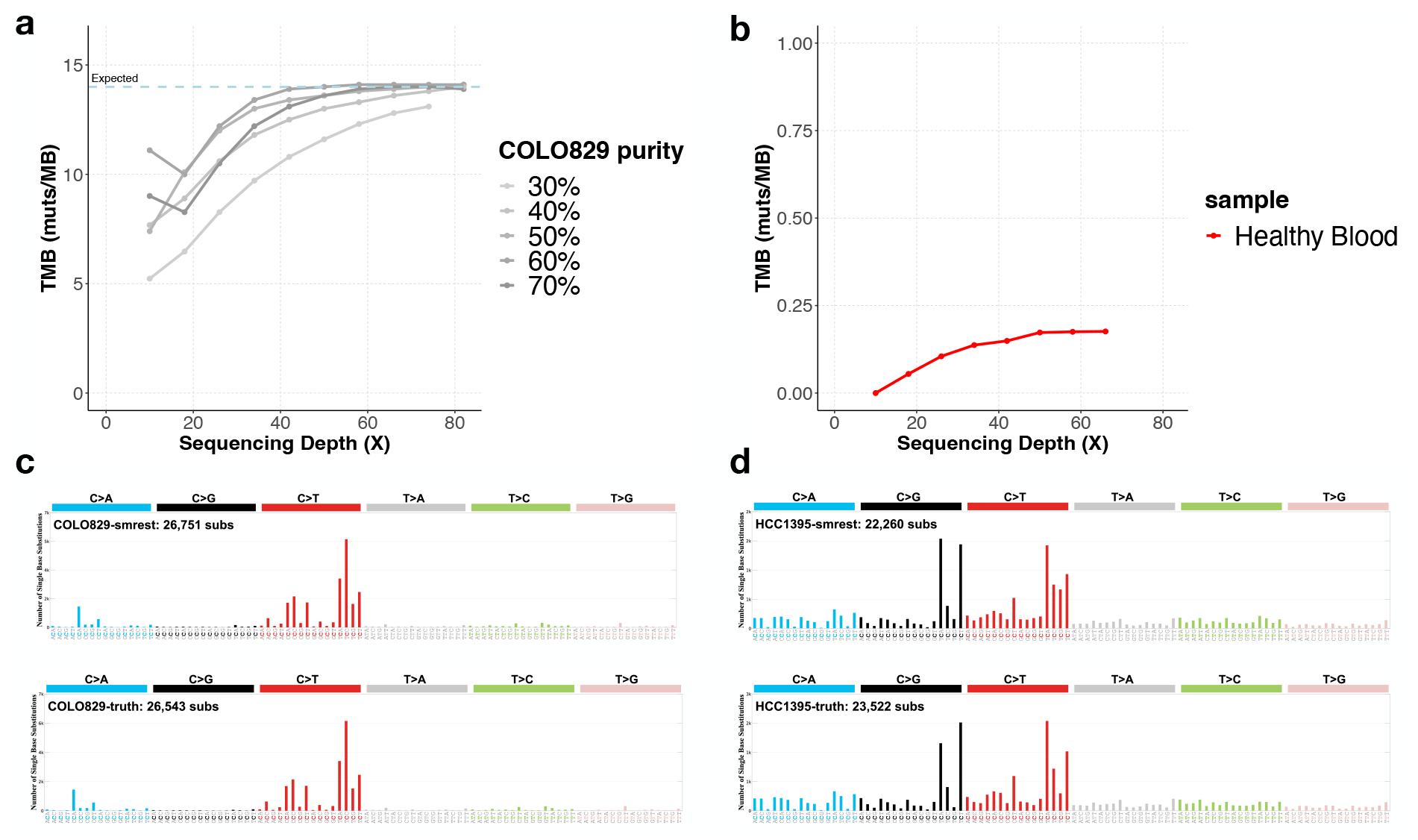
**Panel a** presents the results of estimating tumour mutation burden (TMB) on COLO829 with varying tumour purity (each line series, with darker lines having higher purity) and sequencing depth (x-axis). **Panel b** shows a healthy blood sample (red) as a negative control where few mutations are expected (note different y-axis range than **panel a**). **Panels c** (COLO829) and **d** (HCC1395) show the sequence contexts of called mutations (known as the mutation spectrum) for smrest (top facet) and the ground truth call set (bottom facet) from the 40x/40x experiments.

Mutation signature profiling uses a vector of 96 unique sequence contexts (six different mutation types, each with a single preceding and following base) to determine the relative contribution of mutagenic processes, like UV damage or DNA repair deficiencies (Alexandrov, Nik-Zainal, et al. 2013; Davies et al. 2017). As this analysis requires accurate determination of mutation counts for each sequence context, we assessed the spectrum of mutations found by our program compared to the spectrum of the ground truth data for the 40x/40x datasets for COLO829 (**Figure 5b**) and HCC1395 (**Figure 5c**). The mutation spectrum from smrest is highly similar to that of the ground truth data (cosine similarity *>* 0.99) and predominantly consists of `C>T` mutations in the context `TC>TT` as expected of a sample with UV damage. Similarly, the mutation spectrum for HCC1395 is consistent with the ground truth (cosine similarity *>* 0.98).

## 3 Discussion

In this work we analyzed the potential for calling somatic mutations using single molecule long read sequencing of tumour samples without matched normal samples. Our results suggest that when certain conditions are met - most importantly when tumour purity is within a band determined by sequencing depth - that accurate mutation calling is possible. However, there are limitations to this study that will need to be further explored. The samples sequenced here are all cell lines, where sufficient DNA quantity for long read (and ultra long read) sequencing is easily achieved. Sequencing real tumour samples, particularly solid tumours, will be more challenging and require extensive protocol optimization, or accepting a shorter read length, which will impact the amount of the genome that can be phased and called using this approach. In addition, the cell lines we used are both highly mutated and hence favourable for calculating accuracy statistics given the very large number of true positive mutations found. This approach must also be assessed on tumour samples with a lower mutation rate, although the analysis of the healthy blood sample presented in Figure 5 suggests a low false positive rate.

The algorithms used here are a proof-of-concept that can be improved in a number of ways. Most notably, we do not incorporate allele specific copy number in our mutation classification model unlike in other methods (e.g. Sun et al. 2018). Also, we used a default purity parameter rather than jointly estimating purity and copy number (Carter et al. 2012; Cameron, Baber, Shale, Papenfuss, et al. 2019) as is common in many other approaches. Similarly we do not attempt to infer a cancer cell fraction distribution for subclonal mutations. We use `whatshap` to phase the heterozygous SNPs but this is not designed to account for allele specific copy number changes in cancer, which is an additional source of information. Future work aims to incorporate these improvements, as well as support other mutation types like short indels, and the inference of microsatellite instability, which can be predictive of response to checkpoint inhibitors (K. Li et al. 2020).

## 4 Methods

### 4.1 Simulations

#### 4.1.1 Definitions and notation

- *μ* := somatic mutation rate (probability a given base of the genome is mutated in the tumour)
- *π* := heterozygous SNP rate (probability a given base is a SNP, here fixed at 1*/*1000)
- *α* := tumour purity
- *ϵ* := sequencing error rate
- *a*_*i*_ := number of reads containing the alternate allele at position *i* of the genome
- *r*_*i*_ := number of reads containing the reference allele at position *i* of the genome
- *H*_*j*_[*i*] := the *i*-th base on haplotype *j* (*S*_*j*_[*i*] defined similarly)

#### 4.1.2 Simulated data generation for clonal tumours

For each simulation the average sequencing depth (*λ*) and tumour purity (*α*) are input parameters. To generate the simulated data for a parameter setting (*λ, α*) the following procedure is used. First, the genome size *G* is set to a constant value (here, *G*=100,000,000) then two haplotype strings (*H*_0_, *H*_1_) are initialized from the first *G* bases of human chromosome 2. Each haplotype base was randomly mutated with probability *π* to simulate germline SNPs. A somatic copy of each haplotype (*S*_0_, *S*_1_) was made and mutated at rate *μ*.

For every position *i*, we draw the total sequencing depth *d*_*i*_ from a Poisson(*λ*) distribution and partition the depth across the two haplotypes by drawing *d*_*i*,0_ *∼* Binom(*d*_*i*_, 0.5) and setting *d*_*i*,1_ = *d*_*i*_*−d*_*i*,0_. Then we assign a base to each read. If a position contains a somatic mutation (*S*_*j*_[*i*]*≠H*_*j*_[*i*]) then the base is set to *S*_*j*_[*i*] with probability *α* (the chance of sampling a cancer cell from the mixture) otherwise it is set to *H*_*j*_[*i*]. Finally, the drawn base is randomly changed to one of the three other bases with probability *ϵ* to simulate random sequencing errors. The simulated data is aggregated into a vector containing the number of observed A, C, G and T bases. These vectors were passed into the classifiers described below.

#### 4.1.3 Simulated data classifiers

##### Unphased data

The classifier for unphased data considers three possibilities for each position *i, c*_*i*_ *∈ {*`somatic`, `het`, `reference`*}*, using the number of reads supporting the reference base, denoted *r*_*i*_, and the number of reads supporting the non-reference base with highest read count, *a*_*i*_. It is possible for *a*_*i*_ to be 0 if all reads support the reference base. The classifier for phased data is similar but the classifications and observed data are defined per haplotype *j* (*c*_*i,j*_, *r*_*i,j*_, *a*_*i,j*_).

We calculate the posterior probability a site contains a somatic mutation after observing *a*_*i*_ alternate bases and *r*_*i*_ reference bases as:

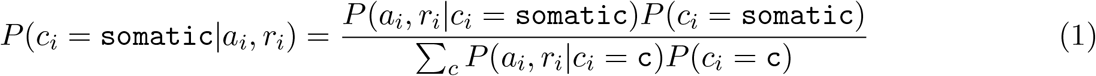

The likelihood term accounts for sequencing errors using a binomial observation model. To observe a read with an alternative base the read must either sample a mutated haplotype (with probability *α/*2) with a correct basecall (1 *− ϵ*), or sample a non-mutated haplotype (with probability 1 *− α/*2) with a base that erroneously supports the alternative allele (*ϵ*). By summing these cases we have the chance of observing an alternative base given the position contains a somatic mutation:

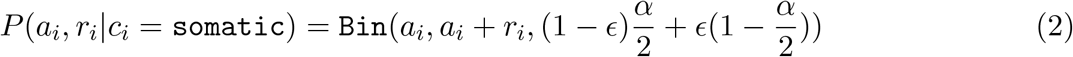

The likelihoods for the heterozygous and reference classifications are similar:

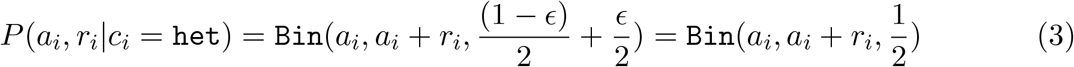

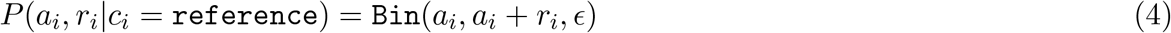

To complete the calculation we use the priors specified by the parameters described above:

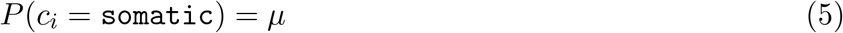

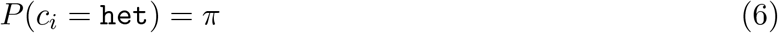

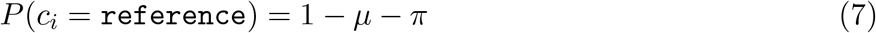

#### 4.1.4 Phased data

When the sequencing data can be phased by assigning each read to a haplotype the problem becomes simpler. To illustrate, consider a position where the reference base is `T` and we have observed 17 reads with `T` and 12 reads with `C`. Under the previous model it is plausible that the position is heterozygous `C/T` and the haplotype bearing the reference allele was sampled more often simply by chance. If the haplotype for each read is known however, we can directly calculate whether the haplotype with the non-reference allele contains sufficient evidence of the reference allele from contaminating non-cancerous cells to classify the position as somatic. Using the example above suppose we have partitioned the reads into two haplotypes where *H*_1_ contains 11 observations of `C` and 5 observations of `T` and *H*_2_ contains 1 observation of `C` and 12 of `T`. It now appears plausible that the position is a `T>C` somatic mutation with the 5 observations of `T` due to contaminating normal cells and the individual’s (inherited) genotype at this position is T/T. Clearly this only works when the sample is not purely tumour (*α <* 1).

The classifier from the previous section can be modified to support phased data:

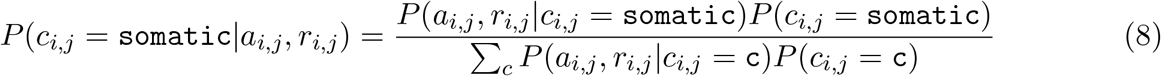

The likelihoods for `somatic` and `reference` classes are similiar to above with the factor of 2 dropped in the `somatic` case (as now the haplotype for each read is known):

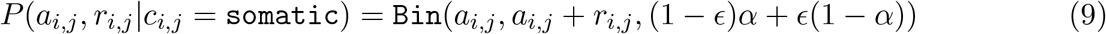

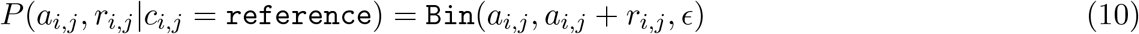

In the case for `het` however, all reads are expected to contain the alternate base, with any reference reads being sequencing errors:

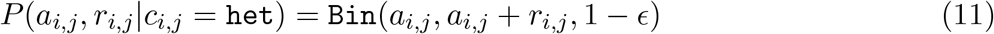

#### 4.1.5 Implementation

The complete simulation procedure is implemented in the function `sim_pileup` in `simulation.rs` in the `smrest` software package.

### 4.2 Single Molecule Long Read Somatic Mutation Calling

The mutation calling procedure for real data is derived from the models presented above. A major difference however is that instead of counting the number of reads supporting the reference and alternate allele, which can be inaccurate if the read-to-reference alignment provided in the BAM file is unreliable, we use a likelihood-based calculation that is common with many other variant callers (Albers et al. 2011; Garrison and Marth 2012; Poplin et al. 2018; Cooke, Wedge, and Lunter 2021). The procedure we use is derived from the Longshot variant caller (Edge and Bansal 2019) with the modification that the core read-haplotype likelihood calculation is changed from Longshot’s Hidden Markov Model to `kprobaln` from HTSlib (H. Li 2011), which incorporates quality scores. In early experiments we found (data not shown) that this change significantly improved somatic mutation calling accuracy.

#### Calculating Read-Haplotype Likelihoods

In the following methods we rely on calculating the probability of observing a certain sequencing read, *R*_*k*_, given a known or assumed haplotype sequence, *H*, denoted *P* (*R*_*k*_|*H*) and referred to as the read-haplotype likelihood. The calculation of *P* (*R*_*k*_|*H*) typically uses the forward algorithm on a hidden Markov model parameterized with gap open, gap extension and substitution probabilities that model the properties of the sequencing platform that generated the read. Longshot also calculates a read-allele likelihood by summing the read-haplotype likelihoods over all haplotypes that contain a particular allele at a particular variant site. We denote this as 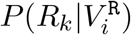and 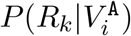 for the reference and alternate alleles of a candidate variant at position *i*. We will denote the set of reads that are informative at site *i* (they cross the reference position and pass all QC filters) as *ℛ*_*i*_.

It is inefficient to perform the forward algorithm over the entire length of each read, so Longshot constrains the calculation to short windows surrounding a position of interest. This procedure involves assembling a set of haplotypes containing combinations of candidate variant alleles, then evaluating *P* (*R*_*k*_|*H*) for each one. We directly use Longshot’s code for performing these calculations and refer to the methods in Edge and Bansal 2019 for further details.

#### Genotyping

The genotyping procedure takes as input a BAM file and a VCF file containing known SNPs in the human population. In the experiments presented in this manuscript we used gnomAD v3 (Karczewski et al. 2020) biallelic SNPs that have allele frequency *>* 0.1%. First, the VCF and BAM file are provided to Longshot’s `extract_fragments` function to calculate read-allele likelihoods. Next we calculate genotype likelihoods for each site in the VCF file. Unlike in typical genotyping applications, where reads at heterozygous sites have equal chance of being drawn from each allele, we account for unbalanced copy number. In the following, let *f* be the probability of drawing a read from the haplotype with higher copy number and *G*_*i*_ be the genotype at position *i*. The genotype likelihoods for the homozygous REF and ALT cases are straightforward:

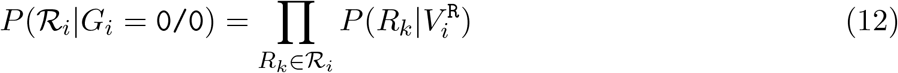

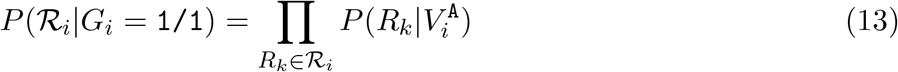

To calculate the heterozygous genotype likelihood we need to account for uncertainty whether the read came from the haplotype bearing the reference or alternate allele, and which one has higher copy number:

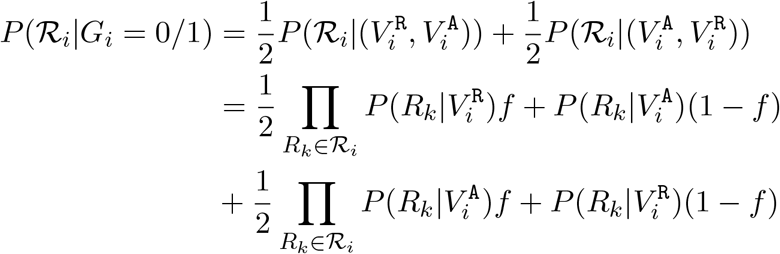

We fix *f* = 0.25 for the experiments in this paper. Future work could fit *f* for each copy number segment of the genome in an iterative procedure that calls an initial set of heterozygous SNPs, estimates *f*, then repeats. Genotype probabilities are calculated using priors that expect approximately 1 in 1,000 sites to be a variant, with 2/3 of variant sites being heterozygous.

The genotype with the highest probability is assigned for each site. Any sites that have evidence of sequencing strand bias (Guo et al. 2012) are left uncalled.

#### Phasing

The VCF file output by the genotyping procedure and the BAM file is provided to `whatshap` (Patterson et al. 2015) for phasing. Default parameters are used, except for the addition of `--ignore-read-groups`. The output is a phased VCF file.

#### Haplotagging

The somatic mutation calling algorithm requires partitioning the input reads by haplotype. Most phasing software, including `whatshap`, can produce a BAM file where reads are annotated with a prediction of which haplotype they originate from, a process known as haplotagging. In smrest we adopt the same procedure, but produce the read-to-haplotype assignments as needed to avoid the time and space required to produce a large BAM file. The output of `whatshap` is set of phased heterozygous SNPs. For simplicity we treat the phased haplotypes as a pair of strings *B*_*j*_ (*j ∈ {*0, 1*}*) where *B*_*j*_[*i*] *∈ {*R, A*}* is the allele at position *i* of haplotype *j*. This formulation neglects the segmentation of the genome into multiple phased blocks, where the phase of adjacent blocks is unknown. This is handled however by our quality control procedure (see below) to discard reads that cross phase block boundaries.

The haplotype assignments can be viewed as a vector *A* where *A*[*k*] *∈ {*0, 1, *−}* is the assignment of read *k* to one of the two haplotypes, or unassigned (*−*). The posterior probability that read *k* originates from haplotype *j* is:

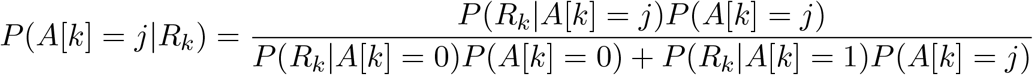

Letting *ℐ*_*k*_ be the set of heterozygous sites that are found in read *k*, the likelihood is computed as the product of the read-variant likelihoods calculated by Longshot:

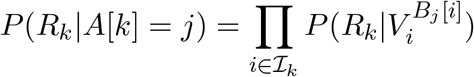

The read is assigned to the most probable haplotype and a quality score (log-scaled probability the haplotype assignment is incorrect) is calculated. If this quality score is less than 20, or if the alleles supported by the read mismatch more than 10% of the alleles in its assigned haplotype, the read is left unassigned and not used for somatic mutation calling. This conservative approach avoids the difficulty of assigning a haplotype to reads that may span multiple phased blocks. Better treatment of these reads is an avenue for future work.

#### Somatic mutation calling

The somatic mutation calling procedure uses the phased VCF file and the original input BAM. For efficiency and parallelization mutations are calculated over 10Mbp windows of the genome. First, reads within the calling window are assigned to a haplotype (if possible) using the procedure described above. Next, a set of candidate somatic variants is found. A position *i* is considered *callable* if both haplotypes have at least 10x sequencing depth and the sum of haplotype depth is not greater than 400x. The callable positions are recorded and output as a BED file. Next, the most frequently observed non-reference base on one of the haplotypes is found. If this base is seen in more than 10% of the reads on the haplotype, and at least 3 times on the haplotype, the position and base are recorded on the list of candidate variants. This procedure is necessarily very permissive and generates a large list of candidate variants, very few of which are expected to be actual somatic mutations. This list of candidate variants is input into Longshot’s `extract_fragments` algorithm as described above to calculate read-allele likelihoods.

Next, every candidate is classified as a somatic mutation, an inherited SNP or reference allele. The general procedure follows the classification described in the simulation section where a reference classification expects all reads to match the reference allele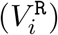, the heterozygous SNP classification expects all reads to match the alternate allele 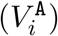 and the somatic mutation expects a mixture of reference and alternative alleles. Here the read-variant likelihoods are used and the model is also extended to account for subclonal mutations. The calculations are performed for each haplotype separately. Letting *R*_*i,j*_ be the set of reads on haplotype *j* that are informative about position *i*:

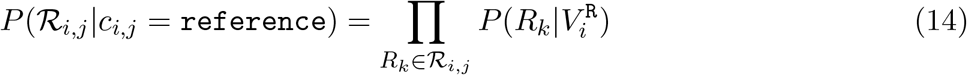

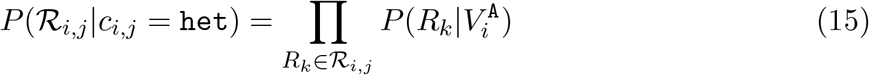

For the somatic class, assuming for the moment all mutations are clonal:

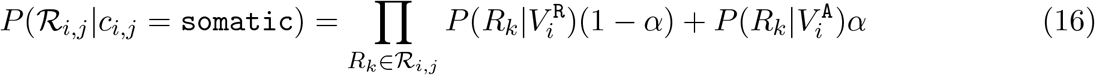

#### Modelling subclonal mutations

The methods described thus far assume that every mutation is clonal and contained in every cancer cell however real tumours have subclonal mutations. We define the *cancer cell fraction*, denoted by *ϕ*_*i*_, as the proportion of tumour cells that carry a mutation at position *i* (for convenience *ϕ*_*i*_ = 0 when position *i* does not have a somatic mutation). For clonal variants *ϕ*_*i*_ = 1, a variant is *subclonal* when 0 *< ϕ*_*i*_ *<* 1. Typically in real cancers a subset of mutations will be clonal, those that arose in the founding lineage of the tumour, so we model the cancer cell fraction distribution as a mixture where a proportion of variants are clonal, denoted *ρ*, and the frequencies for the remaining 1 *− ρ* variants are drawn from a Beta distribution:

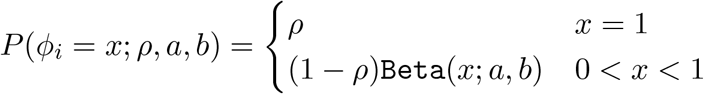

If *ϕ*_*i*_ is known, it would be straightforward to modify the somatic mutation classifier as the chance of sampling a mutated tumour cell is *αϕ*_*i*_. The likelihood then becomes:

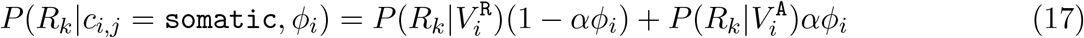

*ϕ*_*i*_ is not known however, so we integrate it out:

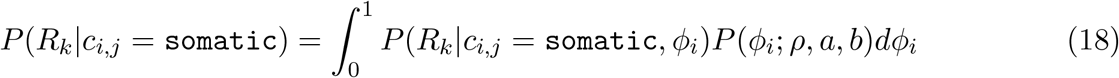

In our experiments we set *ρ* = 0.95, *a* = 2, *b* = 2 for the CCF disribution. The integration is numerically approximated in discrete bins of *ϕ*_*i*_ over the range [0, 1.0].

#### Filtering variants

After identifying somatic mutations we apply filters to remove mutations that either break assumptions of our model or have features that are indicative of problematic regions of the genome. The filters currently used are:

- `MaxOtherHaplotypeObservations`: We expect somatic mutations to appear on only one of the two haplotypes, so use the non-called haplotype as an internal control. If the variant appears on this haplotype more than 2 times, or in more than 20% of the reads, this filter is applied.
- `MinObsPerStrand`: This filter is applied when the variant does not appear in reads from both the forward and reverse sequencing strands, as this is commonly indicative of sequencing artifacts.
- `PossibleAlignmentArtifact`: Our model assumes that the reads used for variant calling (those in *R*_*i,j*_) are reliably mapped and aligned. While Longshot applies QC filters to the reads it uses, primarily mapping quality thresholds, some read alignments may still be erroneous. In particular, we found that reads spanning structural variant breakpoints can have regions with very high mismatch rates, which can be called as somatic mutations, so this filter is applied to remove such calls.
- `LowQual`: the PHRED-scaled mutation quality score falls below the calling threshold.
- `MinHaplotypeDepth`: the depth on either haplotype falls below the calling threshold of 10x coverage.
- `StrandBias`: As in most variant callers we calculate a strand bias p-value to identify possible sequencing artifacts (Guo et al. 2012).

##### Implementation

smrest is implemented in Rust and available under the MIT license on github: https://github.com/jts/smrest. The repository contains a Snakemake (Köster and Rahmann 2018) pipeline that automates the entire process, starting from a BAM file containing reads mapped to the human reference genome.

### 4.3 Experiments

#### Data Access and Preparation

FASTQ, BAM or CRAM files were downloaded from public repositories:

BAM or CRAM files that were already mapped to GRCh38 were used directly, otherwise the reads were mapped with minimap2 or bwa mem for long and short reads, respectively. The raw signal data for the CliveOME data set was downloaded and basecalled with `wf-basecalling` using model `dna_r10.4.1_e8.2_400bps` v4.1.0 in sup mode.

#### Software Versions

The following software tools were used in this work:

#### Downsampling

To prepare BAM files with a specified coverage level we first computed the total number of bases contained in the full depth BAM with `samtools_stats`, calculated the proportion of reads needed to reach the specified coverage, then generated a new BAM by passing this value to the `-s` argument of `samtools_view`.

#### ClairS Mutation Calling

ClairS was run using singularity as described in the README and provided with a tumour and normal BAM file using the `--tumour-bam-fn` and `--normal-bam-fn` arguments. The `--platform` argument was set to `ont_r10_dorado_4khz` for ONT data or `pacbio-hifi` for PacBio-HiFi data.

#### Mutect2 Tumour-Normal Mutation Calling

To generate a raw VCF file the GATK `mutect2` command was run with the panel-of-normals argument set to `1000g_pon.hg38.vcf.gz` and germline resource set to `af-only-gnomad.hg38.vcf.gz`. To force multi-nucleotide variants to be called as individual SNV the `--max-mnp-distance_0` argument was provided. The raw mutation calls were filtered with the `FilterMutectCalls` using the output of the `CalculateContamination` command.

#### Mutect2 Tumour-only Mutation Calling

To call mutations in tumour-only mode, Mutect2 was run as above but provided with a single BAM file containing downsampled reads from both the tumour and normal, and the `--normal-sample-name` argument was omitted.

#### smrest Mutation Calling

smrest was run using the snakemake pipeline implementing the procedure described in the previous section. A single BAM file, containing reads mixed from the tumour and normal cell lines, was provided as input. In all experiments the tumour purity parameter was set to 0.5.

#### Truth Data

The COLO829 truth mutation set was derived from the NovaSeq VCF file provided by the New York Genome Centre (Arora et al. 2019). This VCF file was processed to split MNVs into individual SNVs using `bcftools_norm -a`. The HCC1395 truth mutation set was downloaded from the SEQC2 FTP site.

#### Genome stratification

To evaluate mutation calling performance for all tools, we restricted the analysis to high confidence (HC) regions of the genome. For COLO829 we defined the high confidence regions by first using `bedtools` to compute the union between GIAB’s alldifficultregions and HG001 v4.2.1 complexandSVs BED files (Zook et al. 2014), and ENCODE’s hg38-blacklist.v2 BED (Amemiya, Kundaje, and Boyle 2019). We then took the complement of this union BED file to define the HC regions. For HCC1395 we intersected this BED file with the BED file provided with the truth data from the SEQC FTP.

#### Identifying cell line artifacts

As tumour-only mutation calling may identify mutations acquired in cell culture, which aren’t present in the truth data we used, we removed these calls from our analysis. First, we ran Mutect2 in TN mode but provided the normal sample name in place of the tumour sample name to generate a list of putative mutations in the normal cell line. As loss-of-heterozygosity regions in the tumour would be called as mutations using this procedure, we filtered out any variant call that was within 5,000bp of an annotated population variant (using the `POPAF` field in Mutect2’s VCF) to generate the final list of normal cell line artifacts. Any mutation call matching this list was not included in accuracy calculations.

#### Analyzing Called Mutations

A mutation call set is annotated using the truth data as follows. Each mutation in either the call set or truth set is represented by its chromosome, position, reference allele and alternate allele. The union between the call set and truth set is taken and each record output in a TSV file. Each record is annotated with whether it is contained in the truth set, the call set (excluding hard filtered calls, with the exception of `LowQual` calls as these are retained for calculating precision-recall curves) and within the regions specified within the specified BED file (either the HC regions described above, or the phased subset of these regions). Each record is also annotated with the mutation caller’s confidence (the QUAL field for smrest and clairS, TLOD for Mutect2), the VAF for the truth mutation or called mutation (if applicable) and any filters applied in the VCF file. Accuracy statistics (F1, precision, sensitivity) and precision-recall curves are calculated from this file after removing records where VAF is below the analysis threshold (10%), outside of the regions specified in the BED file or present in the list of normal cell artifacts. When calculating F1, precision or sensitivity a minimum QUAL of 20 was used for smrest calls. Default values were used (`PASS` calls) for clairS and mutect2.

#### Estimating Tumour Mutation Burden

Tumour mutation burden was estimated by counting the number of QC-PASS mutation calls with a minimum QUAL score of 20, and dividing this number by the total number of bases where mutation calling was performed (from the BED file output by smrest). The CliveOME sample listed in **Table 1** was the healthy blood sample control.

#### Mutation Signatures

The mutation spectrum was extracted and plotted using SigProfilerMatrixGenerator (Bergstrom et al. 2019.

#### Implementation

The source code for the mutation caller is provided at https://github.com/jts/smrest. The code used to generate all results in this manuscript is provided as a Snakemake pipeline and associated python scripts at https://github.com/jts/smrest-analysis-pipeline.

## Acknowledgements

The author thanks Joanna Pineda for prior work on somatic mutation calling using phased 10X Genomics Linked Reads (https://github.com/jopineda/10xtrim), and Felix Beaudry and Tom Ouellette for comments on a draft version of this manuscript. The author also thanks Philip Zuzarte, Jim Shaw, Chris Wright, Alvin Ng, Matthew Loose and Winston Timp for discussions related to this manuscript.

The author is supported by the Ontario Institute for Cancer Research through funds provided by the Government of Ontario, the Government of Canada through Genome Canada and Ontario Genomics (OGI-136 and OGI-201) and the National Human Genome Research Institute (NHGRI project 5R01HG009190).

